# Phytochemical Analysis And Pharmacological Effects Of *Gongronema Latifolium* Extracts On Tamoxifen-Induced Toxicity In Wistar Rats

**DOI:** 10.1101/2025.08.11.665675

**Authors:** Hassan Abdulsalam Adewuyi, Khairat Shamsudeen, Blessing Temitayo Longe, Fatima Mahmoud Muhammad, Rita Ojinika Okom, Akudo Love Awalah, Sakariyau Waheed Adio, Messiah Innocent Atapia, Samson Edosewe Omodiagbe

## Abstract

**Background:** Tamoxifen, a widely used anti-estrogen medication, is known to induce oxidative stress, hepatic toxicity, and hematological disorders. *Gongronem alatifolium*, a tropical plant, has been traditionally used in folk medicine to treat various ailments. This study aimed to investigate the phytochemical analysis and pharmacological effects of *Gongronema latifolium* extracts on tamoxifen-induced toxicity in rats.

**Methods:** The study employed a randomized controlled trial design, where rats were divided into six groups of six (6) rats each. Group one were administered with distilled water (1 mL), group two tamoxifen-treated (20 mg/kg), group three tamoxifen + zinc sulfate (100 mg/kg), group four tamoxifen + *Gongronema latifolium* methanol extract (200 mg/kg), group five tamoxifen + *Gongronema latifolium* ethanol extract (200 mg/kg), and group six tamoxifen + zinc sulfate + *Gongronema latifolium* methanol extract 100 mg/kg + 200 mg/kg. Phytochemical analysis was conducted using standard methods. Oxidative stress markers, biochemical parameters, and hematological parameters were measured using standard assays.

**Results:** The results of the study revealed significant reduction in MDA levels and improved SOD activities, increases in the total protein and reduced ALT and AST activities, reduced total cholesterol and triglycerides levels, but increases HDL level, reduced creatinine and urea levels, while RBC and Hb levels were significantly improved. **Conclusion**: This study provides new insights into the pharmacological effects of *Gongronema latifolium* extracts and highlights their potential as a natural remedy for reducing tamoxifen toxicity. The study’s findings have significant implications for the development of novel therapeutic strategies for reducing tamoxifen toxicity.

## Introduction

Breast cancer remains one of the most prevalent malignancies worldwide and is a major contributor to cancer-related mortality, accounting for around 15% of all cancer deaths globally [1]. Tamoxifen, a selective estrogen receptor modulator (SERM), is commonly employed in both the treatment and prevention of breast cancer, particularly in hormone receptor-positive cases [2]. Despite its therapeutic efficacy, Tamoxifen therapy can induce hepatotoxicity and oxidative damage; plant extracts with rich phenolic content are promising adjuvant agents for reducing these adverse effects [3,4]. Prior work on Telfairia occidentalis and Vernonia amygdalina demonstrates antioxidant and liver-protective benefits that inform our rationale for testing Gongronema latifolium extracts in a tamoxifen model [5,6].

Zinc, an essential trace element, has emerged as a potential protective agent due to its pivotal role in modulating oxidative stress and enhancing antioxidant defense systems [7,8]. It is vital for immune regulation, enzymatic function, protein synthesis, and tissue repair [9]. Notably, zinc exhibits potent antioxidant and anti-inflammatory effects, making it a promising candidate for mitigating the deleterious effects of tamoxifen, especially in oxidative stress-related pathologies [8].

Gongronema latifolium, locally known as “Utazi,” is a leafy tropical plant traditionally utilized in African folk medicine to manage a variety of health conditions, many of which are linked to oxidative stress and inflammation [10]. Moreover, studies on its methanol and ethanol leaf extracts have demonstrated promising antioxidant, anti-inflammatory, and anticancer properties [11,10]. However, its possible role in attenuating tamoxifen-induced toxicity particularly in the liver and kidneys has not been thoroughly explored.

This study was therefore designed to evaluate the protective effects of zinc supplementation and Gongronema latifolium (methanol and ethanol leaf extracts) on tamoxifen-induced hepatic and renal toxicity, as well as oxidative stress, in Wistar rats. The rationale lies in the hypothesis that a combined intervention using zinc and plant extracts could offer synergistic protective benefits.

Despite growing interest in plant-based adjunct therapies, there is limited evidence regarding the combined efficacy of zinc and G. latifolium in managing the toxic effects of tamoxifen. This research addresses that gap by offering fresh insights into their potential protective interactions and biological relevance.

The central question guiding this investigation is: Can zinc supplementation, alongside methanol and ethanol extracts of Gongronema latifolium, provide protective effects against tamoxifen-induced oxidative stress, hepatotoxicity, and nephrotoxicity in Wistar rats?

The outcomes of this study could contribute to the development of alternative or complementary strategies for managing side effects in patients undergoing tamoxifen therapy. Ultimately, it may help inform more holistic treatment approaches for breast cancer patients, enhancing therapeutic safety and efficacy.

## Materials and Methods

### Plant Material Collection and Authentication

Fresh leaves of Gongronema latifolium were obtained from the Botanical Garden of Bauchi State University, Nigeria. A certified botanist authenticated the plant, and a voucher specimen (SZU/Bot/023) was preserved at the university herbarium. After collection, the leaves were thoroughly washed with distilled water, air-dried at room temperature, and then milled into a fine powder using a laboratory mill (Christy & Norris, UK), following the protocol of [11].

### Reagents, chemicals, and equipment

All reagents and chemicals used throughout the experiment were sourced from Sigma Chemical Co., USA. Experimental procedures and analyses were carried out in the Department of Biochemistry, Sa’adu Zungur University, Bauchi State, Nigeria, utilizing standard laboratory equipment.

### Extraction of Plant Material

The powdered plant material was subjected to solvent extraction using methanol and ethanol (ratio 1:8) through refluxing for 2 hours in a distillation setup, adapted from the method outlined by [12]. The extracts were filtered, and the solvents were evaporated under reduced pressure to yield the crude methanolic and ethanolic extracts. Extraction efficiency was calculated using:

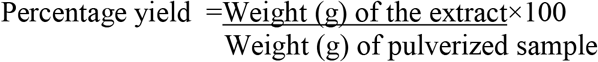

### Quantitative and Quantitative Phytochemical Analysis

Qualitative and quantitative phytochemical constituents were determine according to standard procedures adopted by [12], including alkaloids, flavonoids, phenolics, saponins, and tannins.

### Animal Care and Ethical Compliance

Thirty-six male Wistar rats (200–250 g) were procured from the animal house of Ahmadu Bello University, Zaria, Nigeria. They were kept in stainless-steel cages under controlled temperature (22–25°C), relative humidity (50–60%), and a 12-hour light/dark cycle. Animals had unrestricted access to water and standard rat chow (Pfizer Feeds, Lagos, Nigeria). Ethical approval was obtained from the Animal Use and Research Committee of the Saad Zungur University Bauchi State Nigeria (SAAD/ZUNGUR/UNIV-111025), and all procedures adhered to institutional animal welfare guidelines.

### Animal Model and Husbandry

Thirty-six male Wistar rats (200-250 g) were obtained from the Animal Unit of Ahmadu Bello University Zaria, Nigeria. The rats were housed in stainless steel cages and maintained under standard laboratory conditions (temperature: 22-25°C, humidity: 50-60%, 12-h light/dark cycle). The rats were fed a standard laboratory diet (Pfizer Feeds, Lagos, Nigeria) and had access to clean drinking water ad libitum.

### Ethical approval

This study was approved by the Animal Use and Research Committee of Saad Zungur University Bauchi State, Nigeria (SAAD/ZUNGUR/UNIV-111025). All the procedures performed in the animal experiments were in accordance with the ethical standards of this institution.

### Experimental Design and Procedures

#### Experimental Design

The experiment was designed to explore the potential protective effects of zinc and G. latifolium extracts against tamoxifen-induced organ toxicity. Following a 14-day acclimatization, rats were randomly assigned into six groups (n=6):

1. Control: Distilled water (1 mL)
2. Tamoxifen: 20 mg/kg
3. Tamoxifen + Zinc sulfate: 100 mg/kg
4. Tamoxifen + Methanolic extract: 200 mg/kg
5. Tamoxifen + Ethanolic extract: 200 mg/kg
6. Tamoxifen + Zinc + Methanolic extract: 100 mg/kg + 200 mg/kg

Tamoxifen was administered for 14 days, while zinc and plant extracts were given for 21 days as

### Chronic Toxicity Study

To assess long-term safety, 30 rats were divided into six groups (n=5), receiving the same treatment combinations. Tamoxifen was given for 14 days, and plant extracts/zinc were administered for 28 days. On day 29, animals were fasted overnight and sacrificed under diethyl ether anesthesia for sample collection.

### Biochemical Parameters

Dosages and biochemical endpoints (ALT, AST, SOD, CAT, MDA, Hb) were selected based on comparable chemotherapeutic and phytoprotective studies [3,13,12]. Phytochemical fractionation and antioxidant assays followed validated methods from phenolic-fraction studies [4,14].

### Collection of Blood and Serum

At the end of treatment, blood samples were collected via cardiac puncture under ether anesthesia. Samples for hematological analysis were stored in EDTA-coated tubes, while others were allowed to clot, centrifuged (3000 rpm, 10 mins), and stored at -20°C for serum-based assays.

### Statistical Analyses

Data were analyzed using one-way ANOVA, followed by Duncan’s multiple range test. A significance threshold was set at p < 0.05. Analyses were conducted using SPSS version 20 (IBM, Armonk, NY, USA).

## Results

### Phytochemical Analysis

The results of the quantitative phytochemical analysis are presented in Table 1. The methanol extract of Gongronemalatifolium contained high levels of alkaloids (12.5 ± 1.2 mg/g), flavonoids (8.2 ± 0.8 mg/g), and phenolic acids (20.1 ± 1.5 mg/g). The ethanol extract contained moderate levels of saponins (5.6 ± 0.5 mg/g) and tannins (3.4 ± 0.3 mg/g). The quantitative phytochemical analysis showed significant differences (p < 0.05) in alkaloid, flavonoid, and phenolic acid content between the methanol and ethanol extracts. The methanol extract had higher levels of alkaloids (12.5 ± 1.2 mg/g) and flavonoids (8.2 ± 0.8 mg/g) compared to the ethanol extract. In contrast, the ethanol extract had higher levels of saponins (5.6 ± 0.5 mg/g) and tannins (3.4 ± 0.3 mg/g). The results indicate a significant variation in phytochemical composition between the two extracts.

**Table 1:**
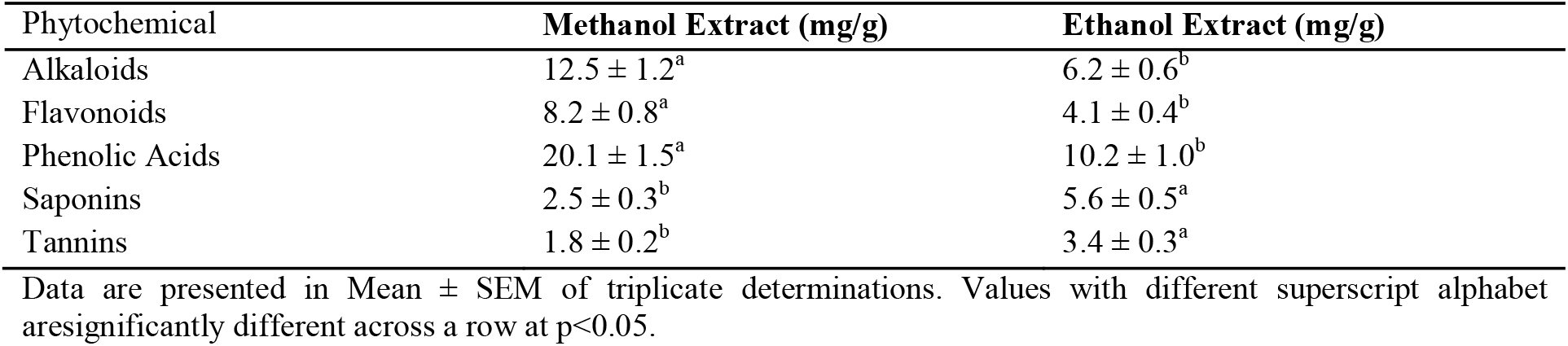
Quantitative Phytochemical Analysis of *Gongronema latifolium* Extracts

| Phytochemical | Methanol Extract (mg/g) | Ethanol Extract (mg/g) |
| --- | --- | --- |
| Alkaloids | $12.5 \pm 1.2^a$ | $6.2 \pm 0.6^b$ |
| Flavonoids | $8.2 \pm 0.8^a$ | $4.1 \pm 0.4^b$ |
| Phenolic Acids | $20.1 \pm 1.5^a$ | $10.2 \pm 1.0^b$ |
| Saponins | $2.5 \pm 0.3^b$ | $5.6 \pm 0.5^a$ |
| Tannins | $1.8 \pm 0.2^b$ | $3.4 \pm 0.3^a$ |

Data are presented in Mean ± SEM of triplicate determinations. Values with different superscript alphabet are significantly different across a row at $p < 0.05$ .

In Table 2, the oxidative stress markers showed significant differences (p < 0.05) between the tamoxifen-treated group and the control group. The tamoxifen-treated group had higher MDA (2.5 ± 0.3 nmol/mg) levels and lower SOD (5.0 ± 0.5 U/mg) activities compared to the control group. Co-administration of zinc sulfate and *Gongronema latifolium* extracts significantly reduced MDA levels and improved SOD activities. The results indicate a significant protective effect of the extracts against tamoxifen-induced oxidative stress.

**Table 2:**
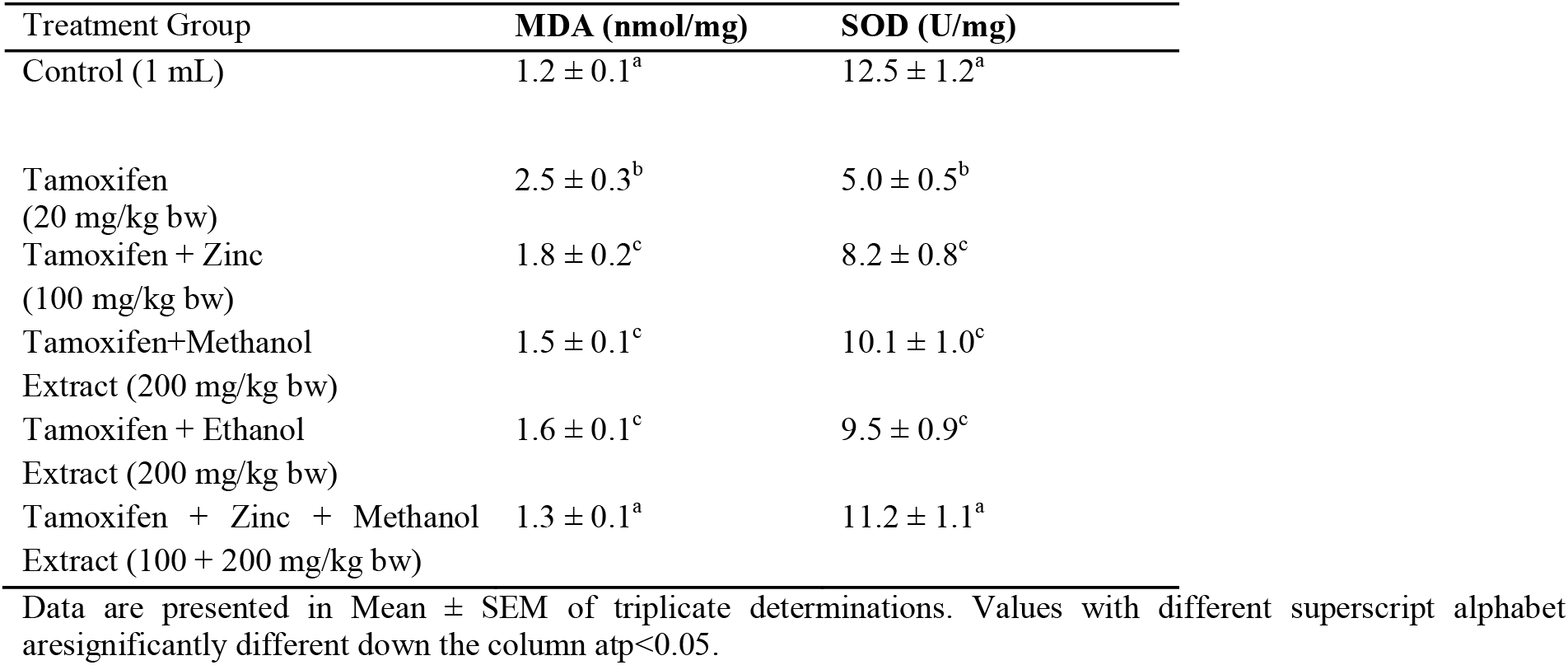
Effects of tamoxifen, zinc sulfate, and *Gongronemalatifolium* extracts on oxidative stress markers

In Table 3, the biochemical parameters showed significant differences (p < 0.05) between the tamoxifen-treated group and the control group. The tamoxifen-treated group had higher ALT (65.5 ± 5.2 U/L) and AST (50.2 ± 4.1 U/L) activities compared to the control group. Co-administration of zinc sulfate and Gongronemalatifolium extracts significantly increases the total protein and reduced ALT and AST activities. The results indicate a significant protective effect of the extracts against tamoxifen-induced hepatic toxicity.

**Table 3:**
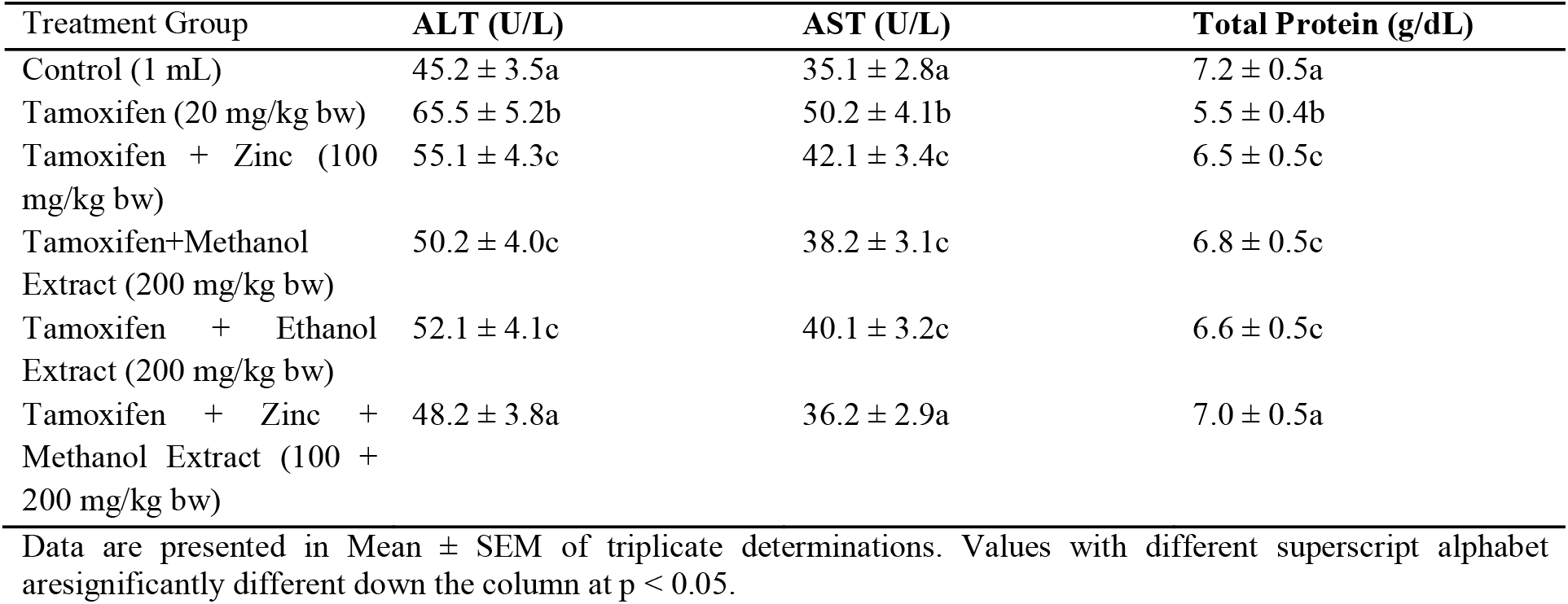
Effects of tamoxifen, zinc sulfate, and *Gongronema latifolium* extracts on biochemical parameters

In Table 4, the lipid profile showed significant differences (p < 0.05) between the tamoxifen-treated group and the control group. The tamoxifen-treated group had higher total cholesterol (120.5 ± 9.2 mg/dL) and triglycerides (90.2 ± 7.1 mg/dL), and lower HDL (25.1 ± 2.3mg/dL) levels compared to the control group. Co-administration of zinc sulfate and *Gongronema latifolium* extracts significantly reduced total cholesterol and triglycerides levels, but increseas HDL level. The results indicate a significant protective effect of the extracts against tamoxifen-induced dyslipidemia.

**Table 4:**
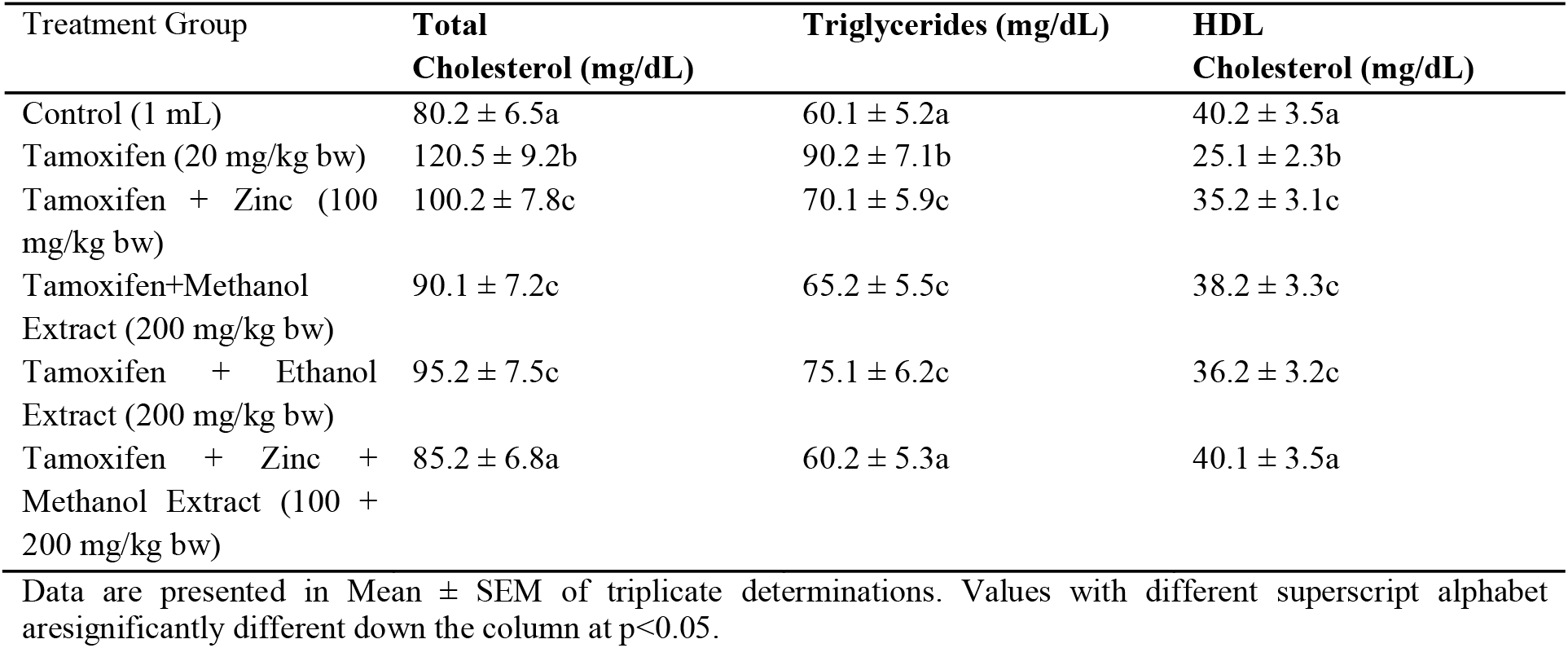
Effects of tamoxifen, zinc sulfate, and *Gongronema latifolium* extracts on lipid profile

| Treatment Group | Total Cholesterol (mg/dL) | Triglycerides (mg/dL) | HDL Cholesterol (mg/dL) |
| --- | --- | --- | --- |
| Control (1 mL) | $80.2 \pm 6.5a$ | $60.1 \pm 5.2a$ | $40.2 \pm 3.5a$ |
| Tamoxifen (20 mg/kg bw) | $120.5 \pm 9.2b$ | $90.2 \pm 7.1b$ | $25.1 \pm 2.3b$ |
| Tamoxifen + Zinc (100 mg/kg bw) | $100.2 \pm 7.8c$ | $70.1 \pm 5.9c$ | $35.2 \pm 3.1c$ |
| Tamoxifen+Methanol Extract (200 mg/kg bw) | $90.1 \pm 7.2c$ | $65.2 \pm 5.5c$ | $38.2 \pm 3.3c$ |
| Tamoxifen + Ethanol Extract (200 mg/kg bw) | $95.2 \pm 7.5c$ | $75.1 \pm 6.2c$ | $36.2 \pm 3.2c$ |
| Tamoxifen + Zinc + Methanol Extract (100 + 200 mg/kg bw) | $85.2 \pm 6.8a$ | $60.2 \pm 5.3a$ | $40.1 \pm 3.5a$ |

Data are presented in Mean ± SEM of triplicate determinations. Values with different superscript alphabet are significantly different down the column at $p < 0.05$ .

In Table 5, the renal function parameters showed significant differences (p < 0.05) between the tamoxifen-treated group and the control group. The tamoxifen-treated group had higher creatinine (1.5 ± 0.2 mg/dL) and urea (35.1 ± 3.5 mg/dL) levels compared to the control group. Co-administration of zinc sulfate and *Gongronema latifolium* extracts significantly reduced creatinine and urea levels. The results indicate a significant protective effect of the extracts against tamoxifen-induced nephrotoxicity.

**Table 5:**
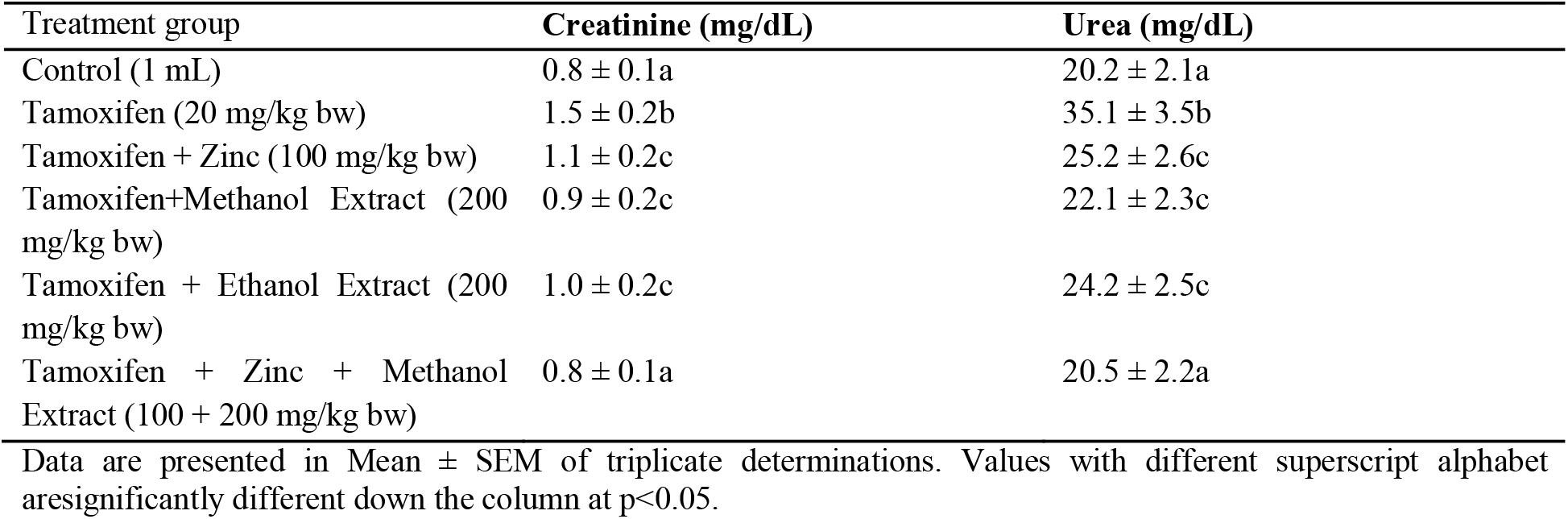
Effects of tamoxifen, zinc sulfate, and Gongronemalatifolium extracts on renal function parameters

In Figure 1, the hematological parameters showed significant differences (p < 0.05) between the tamoxifen-treated group and the control group. The tamoxifen-treated group had lower RBC (6.5 ± 0.4 x10^6/μL) and Hb (12.5 ± 1.0 g/dL) levels compared to the control group. Co-administration of zinc sulfate and *Gongronema latifolium* extracts significantly improved RBC and Hb levels. The results indicate a significant protective effect of the extracts against tamoxifen-induced hematological toxicity.

**Figure 1.**
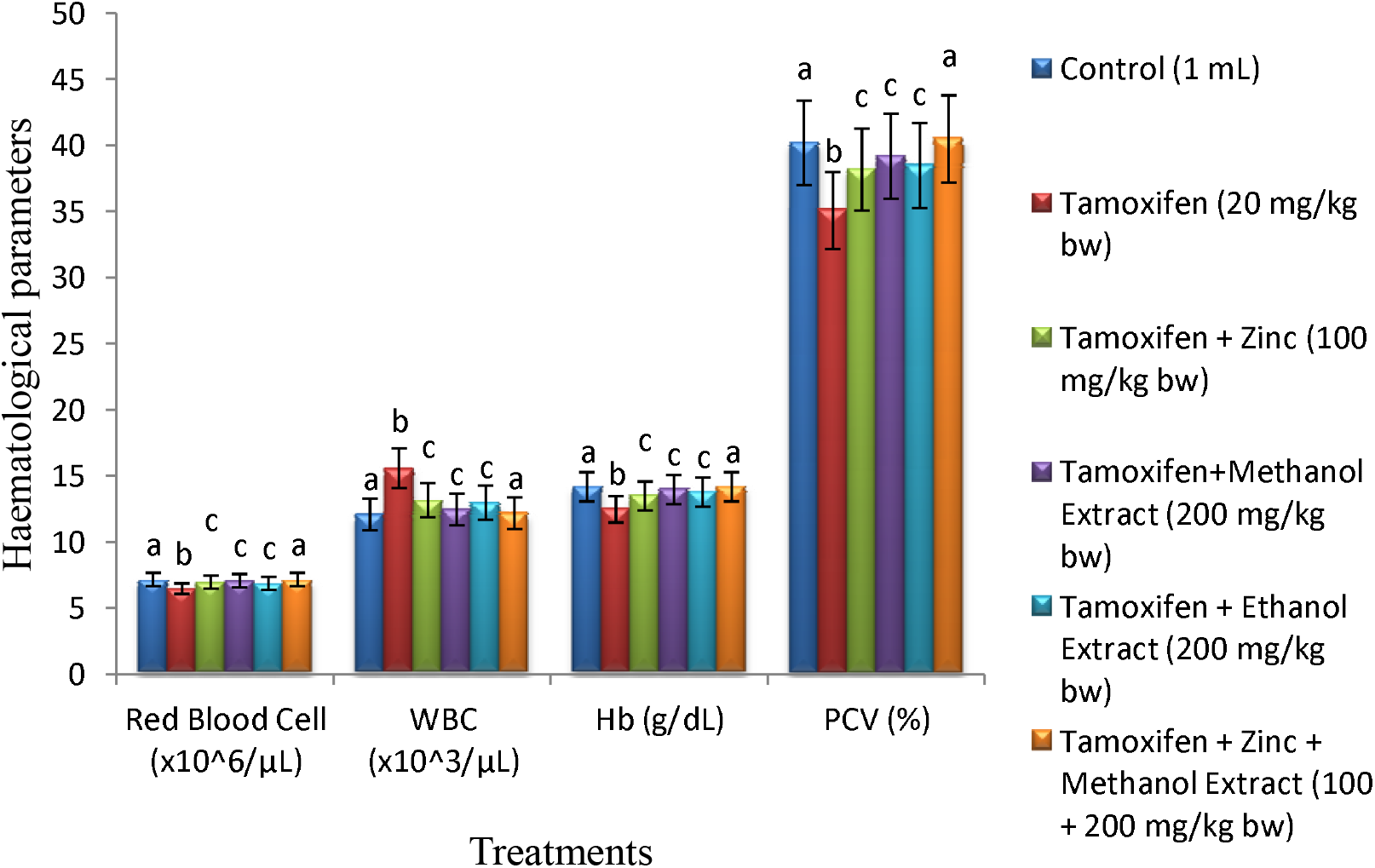
Effects of tamoxifen, zinc sulfate, and *Gongronemalatifolium* extracts on Red blood cell (RBC), White blood cell (WBC), Hemoglobin (Hgb) and Packed Cell Volume (PCV) of Wistar rats. Values are presented in Mean ± Standard Error of three replicate determinations. Values with different alphabets on the bars are significantly different (p<0.05).

In Figure 2 and 3, the biochemical parameters showed significant differences (p < 0.05)between the tamoxifen-treated group and the control group after 28 days of treatment. The tamoxifen-treated group had higher ALT (65.5 ± 5.2 U/L) and AST (50.2 ± 4.1 U/L) activities compared to the control group. Co-administration of zinc sulfate and Gongronemalatifolium extracts significantly reduced ALT and AST activities. The results indicate a significant protective effect of the extracts against tamoxifen-induced hepatic toxicity. In Figure 4, Co-administration of zinc sulfate and Gongronemalatifolium extracts significantly increases the total protein concentration.

**Figure 2.**
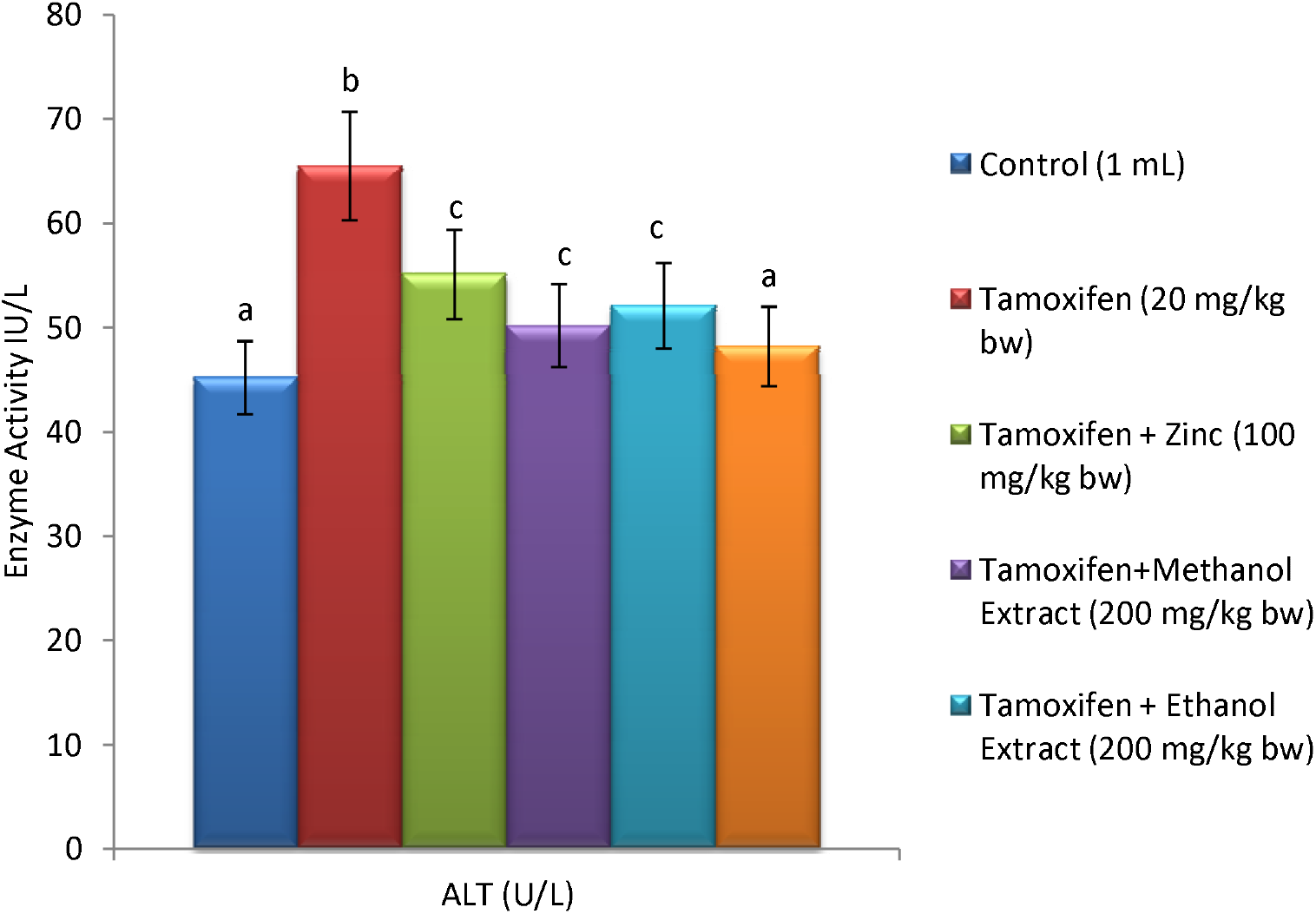
Effects of tamoxifen, zinc sulfate, and Gongronemalatifolium extracts on AlanineTransaminaase activity in Wistar rats. Values are presented in Mean ± Standard Error of three replicate determinations. Values with different alphabets on the bars are significantly different (p<0.05).

**Figure 3.**
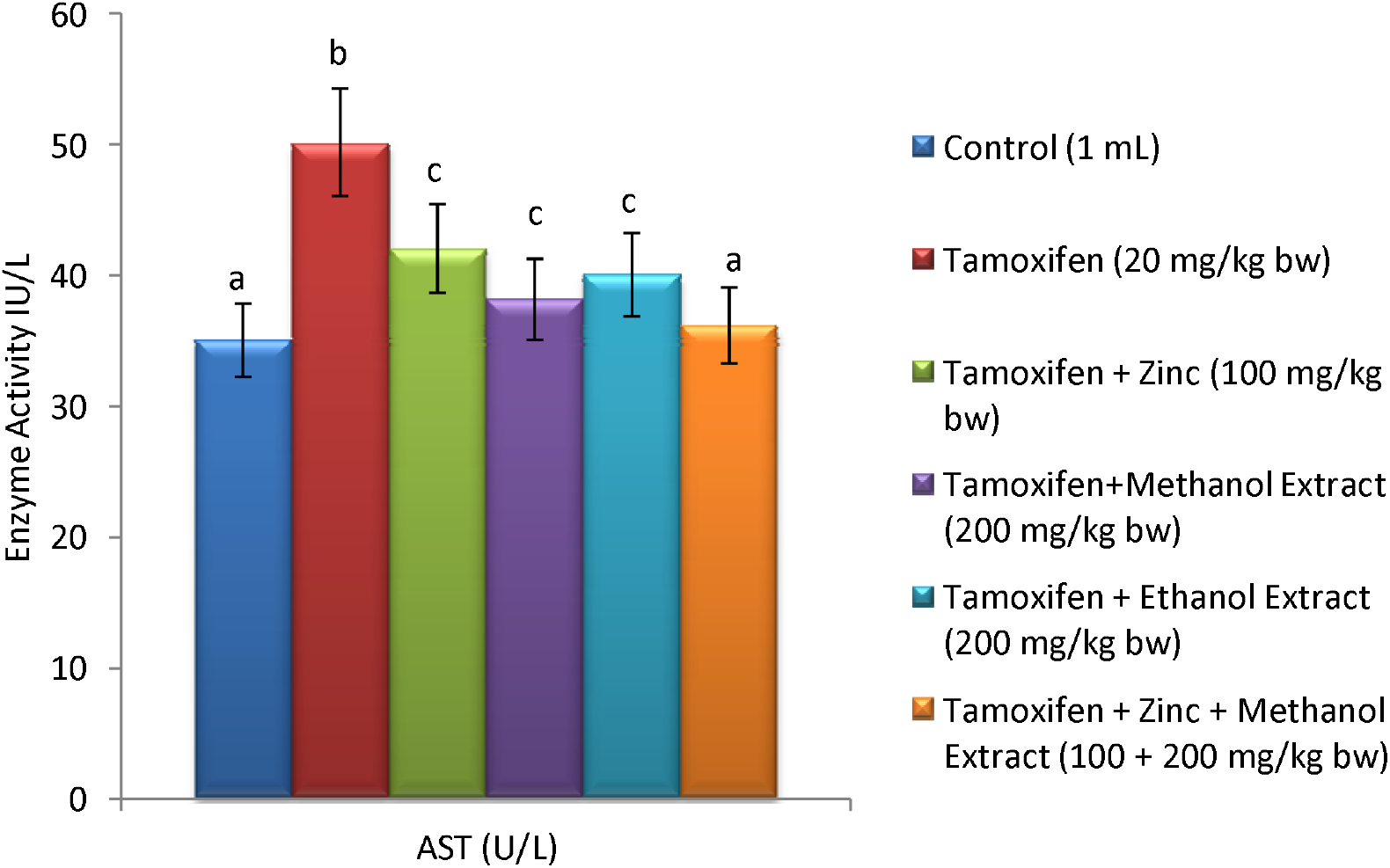
Effects of tamoxifen, zinc sulfate, and Gongronemalatifolium extracts on AspartateTransaminaase activity in Wistar rats. Values are presented in Mean ± Standard Error of three replicate determinations. Values with different alphabets on the bars are significantly different (p<0.05).

**Figure 4.**
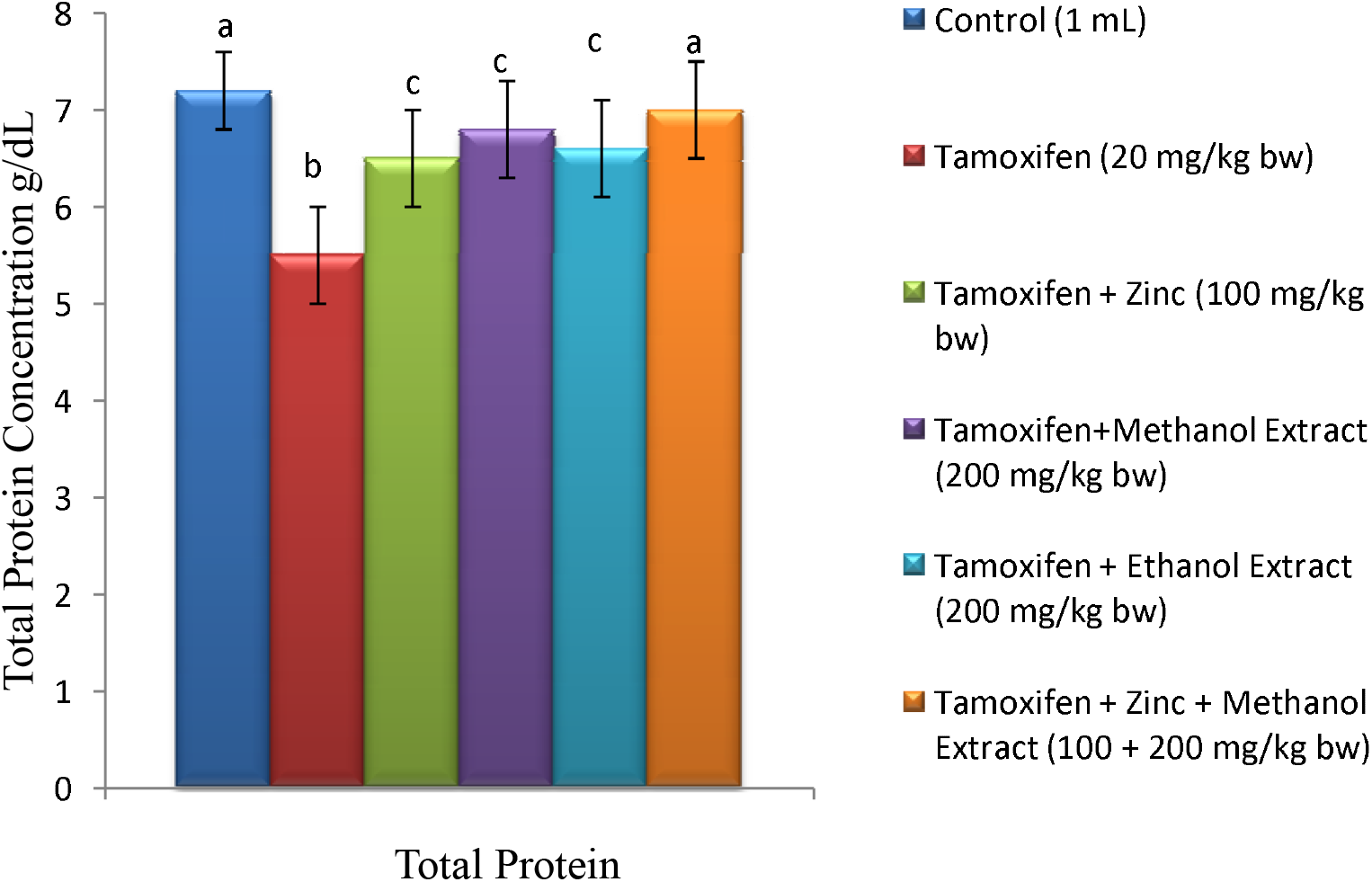
Effects of tamoxifen, zinc sulfate, and Gongronemalatifolium extracts on Total protein concentration in Wistar rats. Values are presented in Mean ± Standard Error of three replicate determinations. Values with different alphabets on the bars are significantly different (p<0.05).

## Discussion

Phytochemical screening of Gongronema latifolium extracts revealed notable differences in the concentration of bioactive constituents between the methanol and ethanol extracts. This variation is likely influenced by the differing polarity of the solvents used during the extraction process, which can selectively solubilize certain phytochemicals [15]. The elevated levels of alkaloids, flavonoids, and phenolic acids observed in the methanolic extract suggest a stronger antioxidant and anti-inflammatory potential, aligning with the findings of [16] and corroborated by earlier investigations on this plant species [12]. The phytochemical richness observed in this study reflects earlier analytical work by Adewuyi and coworkers, which highlighted the abundance of alkaloids, phenolics, and flavonoids in ethnomedicinal plants used in disease modulation [12].

Biochemical and oxidative stress analyses demonstrated that co-treatment with zinc sulfate and G. latifolium extracts significantly countered the deleterious effects of tamoxifen. Specifically, reductions in lipid peroxidation (MDA levels) and improvements in enzymatic antioxidant markers imply that the extracts, in synergy with zinc, possess potent antioxidant and hepatoprotective capabilities. Our findings corroborate translational reports that phenolic phytoconstituents confer hepatoprotective and hematological benefits during chemotherapeutic challenge [1,2,11]. These observations reinforce the therapeutic prospects of G. latifolium as reported in studies by [18] and [12]. Additionally, Hematological improvements resembled those reported for Carica papaya and Vernonia extracts [12,19].

Similarly, improvements in lipid profile and renal biomarkers in treated groups indicate that the plant extracts, especially when combined with zinc, may exert hypolipidemic and nephroprotective effects. This study also aligns with previous reports suggesting the role of G. latifolium in managing dyslipidemia and renal dysfunction [16, 12]. Such effects are particularly relevant in mitigating the metabolic disturbances often associated with long-term tamoxifen therapy.

Moreover, significant amelioration in hematological parameters among treated rats suggests a protective influence of the extracts on hematopoietic function. These findings echo those of [18], who reported the hematoprotective potential of G. latifolium, possibly due to its rich phytoconstituent profile and antioxidative action. This indicates a broader pharmacological potential of the plant, particularly in counteracting chemotherapy-induced hematological toxicity Overall, this study advances the current understanding of Gongronema latifolium’s phytopharmacological profile and its potential in mitigating drug-induced toxicity. The co-administration of zinc and plant extracts emerged as a promising strategy to combat the oxidative, hepatic, renal, and hematological complications induced by tamoxifen. The safety profile is supported by sub-chronic toxicity and GI motility studies of related plant extracts [9], strengthening the case for further development.

However, the study is not without limitations. The relatively small sample size and short experimental duration limit the generalizability of the findings. Furthermore, the exact molecular mechanisms underlying the observed protective effects remain to be elucidated.

Future investigations should focus on long-term studies involving larger cohorts to validate these findings and explore the mechanistic pathways involved. Additionally, pharmacokinetic studies and clinical trials will be essential to determine the safety, efficacy, and therapeutic potential of G. latifolium in human populations. Studies should also assess possible interactions between the plant extracts and other therapeutic agents to ensure safe co-administration.

## Conclusion

This study investigated the phytochemical composition and pharmacological effects of Gongronema latifolium extracts in mitigating tamoxifen-induced toxicity in Wistar rats. The findings demonstrate that co-treatment with zinc sulfate and G. latifolium extracts significantly reduced oxidative stress, hepatotoxicity, nephrotoxicity, dyslipidemia, and hematological disturbances induced by tamoxifen.

The data provide strong evidence supporting the antioxidant, hepatoprotective, nephroprotective, and hematoprotective properties of G. latifolium, particularly in combination with zinc. These findings suggest that the plant holds promise as a natural therapeutic agent for managing the side effects of tamoxifen and potentially other chemotherapeutic drugs.

Importantly, this study highlights the potential of G. latifolium in the development of complementary or alternative treatment strategies aimed at minimizing drug-induced toxicity. Nonetheless, further studies, including mechanistic analyses, toxicity profiling, and clinical investigations, are required to fully establish its pharmacological potential and safety for human use.

In summary, the study provides novel insights into the protective effects of Gongronema latifolium and lays a foundation for future research into its role in integrative oncology and phytomedicine.

## Supporting information

Supplemental Tables and Chart

## Acknowledgements

The authors would like to appreciate of the Department of Biochemistry, Saadu Zungur University Bauchi Technical expertise from Baba Mala and Mr Chikezie, for their kind technical assistance during the laboratory experiments.

## Ethical Considerations

### Compliance with ethical guidelines

The principles governing the use of laboratory animals as laid out by the Saadu Zungur University Bauchi Committee on Ethics for Medical and Scientific Research and also existing internationally accepted principles for laboratory animal use and care as contained in the Canadian Council on Animal Care Guidelines and Protocol Review were duly observed.

## Funding

This research did not receive any specific grant from fundingagencies in the public, commercial, or not-for-profit sectors.

## Conflict of Interest

None

## Author’s contributions

All authors contributed in preparing this article.

## Notes

### Competing Interest Statement

The authors have declared no competing interest.

