## Supplemental Tables and Chart for "Phytochemical Analysis And Pharmacological Effects Of *Gongronema Latifolium* Extracts On Tamoxifen-Induced Toxicity In Wistar Rats"

**Table 1: Quantitative Phytochemical Analysis of *Gongronemalatifolium* Extracts**

| Phytochemical | Methanol Extract (mg/g) | Ethanol Extract (mg/g) |
| --- | --- | --- |
| Alkaloids | 12.5 ± 1.2^a^ | 6.2 ± 0.6^b^ |
| Flavonoids | 8.2 ± 0.8^a^ | 4.1 ± 0.4^b^ |
| Phenolic Acids | 20.1 ± 1.5^a^ | 10.2 ± 1.0^b^ |
| Saponins | 2.5 ± 0.3^b^ | 5.6 ± 0.5^a^ |
| Tannins | 1.8 ± 0.2^b^ | 3.4 ± 0.3^a^ |

Data are presented in Mean ± SEM of triplicate determinations. Values with different superscript alphabet aresignificantly different across a row at p<0.05.

**Table 2: Effects of tamoxifen, zinc sulfate, and *Gongronemalatifolium* extracts on oxidative stress markers**

| Treatment Group | MDA (nmol/mg) | SOD (U/mg) |
| --- | --- | --- |
| Control (1 mL) | 1.2 ± 0.1^a^ | 12.5 ± 1.2^a^ |
| Tamoxifen  (20 mg/kg bw) | 2.5 ± 0.3^b^ | 5.0 ± 0.5^b^ |
| Tamoxifen + Zinc  (100 mg/kg bw) | 1.8 ± 0.2^c^ | 8.2 ± 0.8^c^ |
| Tamoxifen+Methanol  Extract (200 mg/kg bw) | 1.5 ± 0.1^c^ | 10.1 ± 1.0^c^ |
| Tamoxifen + Ethanol  Extract (200 mg/kg bw) | 1.6 ± 0.1^c^ | 9.5 ± 0.9^c^ |
| Tamoxifen + Zinc + Methanol Extract (100 + 200 mg/kg bw) | 1.3 ± 0.1^a^ | 11.2 ± 1.1^a^ |

Data are presented in Mean ± SEM of triplicate determinations. Values with different superscript alphabet aresignificantly different down the column atp<0.05.

Table 3: Effects of tamoxifen, zinc sulfate, and *Gongronemalatifolium* extracts on biochemical parameters

| Treatment Group | ALT (U/L) | AST (U/L) | Total Protein (g/dL) |
| --- | --- | --- | --- |
| Control (1 mL) | 45.2 ± 3.5a | 35.1 ± 2.8a | 7.2 ± 0.5a |
| Tamoxifen (20 mg/kg bw) | 65.5 ± 5.2b | 50.2 ± 4.1b | 5.5 ± 0.4b |
| Tamoxifen + Zinc (100 mg/kg bw) | 55.1 ± 4.3c | 42.1 ± 3.4c | 6.5 ± 0.5c |
| Tamoxifen+Methanol Extract (200 mg/kg bw) | 50.2 ± 4.0c | 38.2 ± 3.1c | 6.8 ± 0.5c |
| Tamoxifen + Ethanol Extract (200 mg/kg bw) | 52.1 ± 4.1c | 40.1 ± 3.2c | 6.6 ± 0.5c |
| Tamoxifen + Zinc + Methanol Extract (100 + 200 mg/kg bw) | 48.2 ± 3.8a | 36.2 ± 2.9a | 7.0 ± 0.5a |

Data are presented in Mean ± SEM of triplicate determinations. Values with different superscript alphabet aresignificantly different down the column at p<0.05.

Table 4: Effects of tamoxifen, zinc sulfate, and Gongronemalatifolium extracts on lipid profile

| Treatment Group | Total  Cholesterol (mg/dL) | Triglycerides (mg/dL) | HDL  Cholesterol (mg/dL) |
| --- | --- | --- | --- |
| Control (1 mL) | 80.2 ± 6.5a | 60.1 ± 5.2a | 40.2 ± 3.5a |
| Tamoxifen (20 mg/kg bw) | 120.5 ± 9.2b | 90.2 ± 7.1b | 25.1 ± 2.3b |
| Tamoxifen + Zinc (100 mg/kg bw) | 100.2 ± 7.8c | 70.1 ± 5.9c | 35.2 ± 3.1c |
| Tamoxifen+Methanol Extract (200 mg/kg bw) | 90.1 ± 7.2c | 65.2 ± 5.5c | 38.2 ± 3.3c |
| Tamoxifen + Ethanol Extract (200 mg/kg bw) | 95.2 ± 7.5c | 75.1 ± 6.2c | 36.2 ± 3.2c |
| Tamoxifen + Zinc + Methanol Extract (100 + 200 mg/kg bw) | 85.2 ± 6.8a | 60.2 ± 5.3a | 40.1 ± 3.5a |

Data are presented in Mean ± SEM of triplicate determinations. Values with different superscript alphabet aresignificantly different down the column at p<0.05.

Table 5: Effects of tamoxifen, zinc sulfate, and Gongronemalatifolium extracts on renal function parameters

| Treatment group | Creatinine (mg/dL) | Urea (mg/dL) |
| --- | --- | --- |
| Control (1 mL) | 0.8 ± 0.1a | 20.2 ± 2.1a |
| Tamoxifen (20 mg/kg bw) | 1.5 ± 0.2b | 35.1 ± 3.5b |
| Tamoxifen + Zinc (100 mg/kg bw) | 1.1 ± 0.2c | 25.2 ± 2.6c |
| Tamoxifen+Methanol Extract (200 mg/kg bw) | 0.9 ± 0.2c | 22.1 ± 2.3c |
| Tamoxifen + Ethanol Extract (200 mg/kg bw) | 1.0 ± 0.2c | 24.2 ± 2.5c |
| Tamoxifen + Zinc + Methanol Extract (100 + 200 mg/kg bw) | 0.8 ± 0.1a | 20.5 ± 2.2a |

Data are presented in Mean ± SEM of triplicate determinations. Values with different superscript alphabet aresignificantly different down the column at p<0.05.

Haematological parameters

| Treatments | Control (1 mL) | Tamoxifen (20 mg/kg bw) | Tamoxifen + Zinc (100 mg/kg bw) | Tamoxifen+Methanol Extract (200 mg/kg bw) | Tamoxifen + Ethanol Extract (200 mg/kg bw) | Tamoxifen + Zinc + Methanol Extract (100 + 200 mg/kg bw) |
| --- | --- | --- | --- | --- | --- | --- |
| Red Blood Cell (x10^6/μL) | 7.2 | 6.5 | 7 | 7.1 | 6.9 | 7.2 |
| WBC (x10^3/μL) | 12.1 | 15.6 | 13.2 | 12.5 | 13 | 12.2 |
| Hb (g/dL) | 14.2 | 12.5 | 13.5 | 14 | 13.8 | 14.2 |
| PCV (%) | 40.2 | 35.1 | 38.2 | 39.2 | 38.5 | 40.5 |

**Figure 1.**Effects of tamoxifen, zinc sulfate, and *Gongronemalatifolium* extracts on Red blood cell (RBC), White blood cell (WBC), Hemoglobin (Hgb) and Packed Cell Volume (PCV) of Wistar rats

Enzyme Activity IU/L

|  | Control (1 mL) | Tamoxifen (20 mg/kg bw) | Tamoxifen + Zinc (100 mg/kg bw) | Tamoxifen+Methanol Extract (200 mg/kg bw) | Tamoxifen + Ethanol Extract (200 mg/kg bw) | Tamoxifen + Zinc + Methanol Extract (100 + 200 mg/kg bw) |
| --- | --- | --- | --- | --- | --- | --- |
| ALT (U/L) | 45.2 | 65.5 | 55.1 | 50.2 | 52.1 | 48.2 |

**Figure 2.**Effects of tamoxifen, zinc sulfate, and Gongronemalatifolium extracts on AlanineTransaminaase activity in Wistar rats

Enzyme Activity IU/L

|  | Control (1 mL) | Tamoxifen (20 mg/kg bw) | Tamoxifen + Zinc (100 mg/kg bw) | Tamoxifen+Methanol Extract (200 mg/kg bw) | Tamoxifen + Ethanol Extract (200 mg/kg bw) | Tamoxifen + Zinc + Methanol Extract (100 + 200 mg/kg bw) |
| --- | --- | --- | --- | --- | --- | --- |
| AST (U/L) | 35.1 | 50.2 | 42.1 | 38.2 | 40.1 | 36.2 |

**Figure 3.**Effects of tamoxifen, zinc sulfate, and Gongronemalatifolium extracts on AspartateTransaminaase activity in Wistar rats

Values are presented in Mean ± Standard Error of three replicate determinations. Values with different alphabets on the bars are significantly different (p<0.05).

Total Protein Concentration g/dL

| Total Protein | Control (1 mL) | Tamoxifen (20 mg/kg bw) | Tamoxifen + Zinc (100 mg/kg bw) | Tamoxifen+Methanol Extract (200 mg/kg bw) | Tamoxifen + Ethanol Extract (200 mg/kg bw) | Tamoxifen + Zinc + Methanol Extract (100 + 200 mg/kg bw) |
| --- | --- | --- | --- | --- | --- | --- |
|  | 7.2 | 5.5 | 6.5 | 6.8 | 6.6 | 7 |

**Figure 4.**Effects of tamoxifen, zinc sulfate, and Gongronemalatifolium extracts on Total protein concentration in Wistar rats

Values are presented in Mean ± Standard Error of three replicate determinations. Values with different alphabets on the bars are significantly different (p<0.05).
